# Detection of breast cancer lymph node metastases in frozen sections with a point-of-care low-cost microscope scanner

**DOI:** 10.1101/474106

**Authors:** Oscar Holmström, Nina Linder, Hannu Moilanen, Antti Suutala, Stig Nordling, Anders Ståhls, Mikael Lundin, Vinod Diwan, Johan Lundin

## Abstract

**Background:** Detection of lymph node metastases is essential in breast cancer diagnostics and staging, affecting treatment and prognosis. Intraoperative microscopy analysis of sentinel lymph node frozen sections is standard for detection of axillary metastases, but requires access to a pathologist for sample analysis. Remote analysis of digitized samples is an alternative solution, but is limited by the requirement for high-end slide scanning equipment.

**Objective:** To determine whether the image quality achievable with a low-cost, miniature digital microscope scanner is sufficient for detection of metastases in breast cancer lymph node frozen sections.

**Methods:** Lymph node frozen sections from 79 breast cancer patients were digitized using a prototype miniature microscope scanner and a high-end slide scanner. Images were independently reviewed by two pathologists and results compared between devices with conventional light microscopy analysis as ground truth.

**Results:** Detection of metastases in the images acquired with the miniature scanner yielded an overall sensitivity of 91 % and specificity of 99 % and showed strong agreement when compared to light microscopy (*k* = 0.91). Strong agreement was also observed when results were compared to results from the high-end slide scanner (*k* = 0.94). A majority of discrepant cases were micrometastases and sections of which no anticytokeratin staining was available.

**Conclusion:** Accuracy of detection of metastatic cells in breast cancer sentinel lymph node frozen sections by visual analysis of samples digitized using low-cost, point-of-care microscopy is comparable to analysis of digital samples scanned using a high-end, whole slide scanner. This technique could potentially provide a workflow for digital diagnostics in resource-limited settings, facilitate sample analysis at the point-of-care and reduce the need for trained experts on-site during surgical procedures.

## Introduction

Breast cancer is the most common form of cancer in women, and the second leading cause of cancer-related death in women globally [1]. Detection of axillary lymph node metastases remains essential for the staging of breast cancer, affecting treatment and prognosis [2]. Presence of axillary lymph node metastases indicates a need for more extensive surgical procedures, typically axillary lymph node dissection (ALND) [3]. Axillary metastases can be detected accurately using sentinel lymph node biopsies in the vast majority of node positive patients, thus avoiding unnecessary further axillary surgery for node negative patients [4, 5]. This is important as evacuation of axillary lymph nodes is a major cause of postoperative complications [6]. Intraoperative evaluation of frozen sections from sentinel lymph nodes (FS) is the most common technique to determine axillary lymph node status, but requires the presence of a pathologist on-site or close to the point-of-care to analyze the samples. Surgical pathology using FS is generally considered accurate for the detection of macrometastases, but not as reliable for detection of smaller lesions, i.e. micrometastases and isolated tumor cells [7, 8]. Light microscopy evaluation of FS is also prone to a certain degree of subjectivity [9].

During the last decade, the field of digital pathology has evolved significantly. Whole-slide imaging (i.e. slide digitization) is now feasible with magnification and spatial image quality comparable to conventional light microscopy [10]. Digital pathology using digitized microscopy samples, or whole slide images (WSI), has multiple advantages, such as enabling remote access to samples for consultation purposes and remote sample analysis, and thus reducing the need for on-site experts. Another significant advantage is the possibility of utilizing digital image analysis to facilitate sample analysis [11]. Studies suggest that the use of WSI to interpret FS samples at a distance is feasible with results comparable to conventional methods [12], and this technique is already being utilized in clinical settings at certain locations where on-site access to a pathologist is limited [13]. Currently however, the digitization of FS has to be carried out with high-end, whole slide scanners, which mainly due to their high cost (retail prices ranging from 30 000 - 200 000 €) are limited to well-equipped clinics. These devices also tend to be bulky in size and require trained personnel and regular maintenance, further limiting their usability for point-of-care slide scanning [14].

During recent years, studies have demonstrated how extremely cost-efficient digital microscopy devices for point-of-care microscopy diagnostics can be constructed using commonly available, low-cost, mass-produced components from consumer electronic products (typically smart phone camera systems) [15]. As the performance of smart phone cameras has improved significantly during the last decade, the imaging performance of this type of devices has also increased accordingly. Studies suggest that the image quality achievable with this type of devices and components is sufficient for diagnostic purposes in a variety of diseases, such as parasitic diseases [16, 17], routine cancer histopathology [18]. Currently these devices have certain limitations, one being that the digitized area typically is limited to a single field-of-view (FOV) of the device.

Here, we studied the performance of a prototype, low-cost, mobile digital microscope scanner which supports digitization of sample areas measuring multiple FOVs, i.e. whole slide imaging. We evaluate the performance of the device for digitization of routinely prepared, intraoperative breast cancer frozen sections. The WSIs captured with the miniature microscope prototype are assessed by two independent pathologists to detect metastases and results compared to conventional microscopy and to analysis of WSIs captured with a high-end scanner.

## Materials and methods

### Sample collection

Samples used in this study were routinely collected sentinel lymph node frozen sections, acquired during breast cancer surgery at hospitals within the Hospital District of Helsinki and Uusimaa in southern Finland. The samples were collected and prepared in accordance with local standard operating procedures during a period of one year (2016), and archived in the files of the pathology laboratory of the hospital district (HUSLAB, Helsinki, Finland). Frozen sections were cut with a thickness of 5 µm, and routine staining performed using toluidine blue and anti-cytokeratin immunohistochemical staining. Immunostaining for cytokeratins was performed with a staining kit containing mouse monoclonal antibodies, targeting a variety of cytokeratins, and diamino benzidine as a chromogen (Cytonel Plus kit, Jilab Inc., Tampere, Finland).

For this study, we retrospectively identified and collected samples from a total number of 80 patients. Of these, 28 patients were node positive (i.e. histologically verified macro- or micrometastases) and 52 patients were node negative (i.e. no detected cancer cells). A majority of patients had sections stained with both toluidine blue and anti-cytokeratin antibodies, but for a minority of selected patients only toluidine blue sections (n = 3) or anti-cytokeratin stained sections (n = 3) were available. For this study, we decided to limit the analysis to one area of representative tissue from every glass slide, selected visually by light microscopy expert evaluation. For every patient, one representative glass slide stained with toluidine blue and the corresponding slide, stained with anti-cytokeratin (if available) was collected after which representative sample regions, measuring approximately 0.5 x 0.5 cm (25 mm^2^), were selected and marked by a pathologist (SN) for digitization and further analysis.

The ground truth in the study was decided as the light microscopy diagnosis of the physical frozen sections, performed by a pathologist experienced in breast cancer pathology. Thus, after the slides had been collected, all slides were examined by a pathologist (SN) to confirm diagnosis used as the study ground truth. One sample was excluded during this phase, as staining artefacts affected sample quality.

### Digitization of samples

The evaluated instrument is a portable, lightweight, cloud-connected digital microscope scanner prototype developed by the Institute for Molecular Medicine Finland – FIMM, University of Helsinki (Fig 1). The imaging optics of the microscope is constructed using low-cost, mass produced polymer lenses, primarily developed for usage in cell phone camera systems. The prototype was manufactured by a company specialized in providing services for the microelectronics industry (Laser Probe LP Oy, Oulu, Finland). Total material costs for the miniaturized imaging optics in the device, including the integrated focusing system, are comparable to costs of the optics of a mid-range smartphone. A white light-emitting diode (LED) is used as the source of light for brightfield imaging, and by utilizing a retractable ultraviolet LED source with adjacent filters, transmitted light fluorescence imaging is also supported. The camera module (See3CAM_130, e-con Systems Inc., St Louis, USA) of the microscope features a 13-megapixel complementary metal oxide semiconductor (CMOS) sensor with a plastic 1/3.2” lens and a maximum image resolution of 4208 x 3120 pixels. The field of view of the microscope is approximately 0.93 x 0.69 mm^2^ with a pixel size of approximately 0.22 µm x 0.22 µm and the spatial resolution 0.9 µm, as measured using a standardized USAF resolution test chart (Fig 2). Coarse focus is adjusted manually using a physical focus lever to adjust focus plane, and fine focus automatically using the built-in auto focus-routine. The device is connected, powered and operated through a universal serial bus (USB) connector from a computer running a custom software written in the matrix laboratory programming environment (MATLAB, MathWorks Inc, Natick, MA) to control the device. The software features a live-view of the sample area, and controls to select and adjust areas to be scanned. Adjustment of the glass slide can be performed manually, or by utilizing the external motor unit to adjust sample position. Digitization of larger sample areas (i.e. whole slide scanning), covering multiple field of views, is possible by utilizing the external motor unit for automatic sample navigation while the device automatically captures a series of images from the different location. Acquired images are saved locally on the computer and uploaded to an image processing and management platform (WebMicroscope, Fimmic Oy, Helsinki, Finland) running on a cloud server located at the university campus (Meilahti Campus Library Terkko, University of Helsinki, Helsinki, Finland). Scanned areas measuring multiple FOVs are stitched together after the scanning process into a single virtual slide. We used the commercially available software Image Composite Editor (Microsoft Computational Photography Research Group, Microsoft Inc., Redmond, WA) for the image stitching process. The generated digital samples were saved in the Tagged Image File Format (TIFF), and further compressed to a wavelet file format (Enhanced Compressed Wavelet; ECW, Hexagon Geospatial, Wisconson, USA) with a target compression ratio of 1:9 to reduce file size, before uploading to the image management platform. As shown in earlier work, this amount of compression preserves sufficient spatial detail to not alter results significantly [19]. Remote access to the image server for sample viewing and scoring was established using a web browser, secured with SSL encryption.

**Fig 1.**
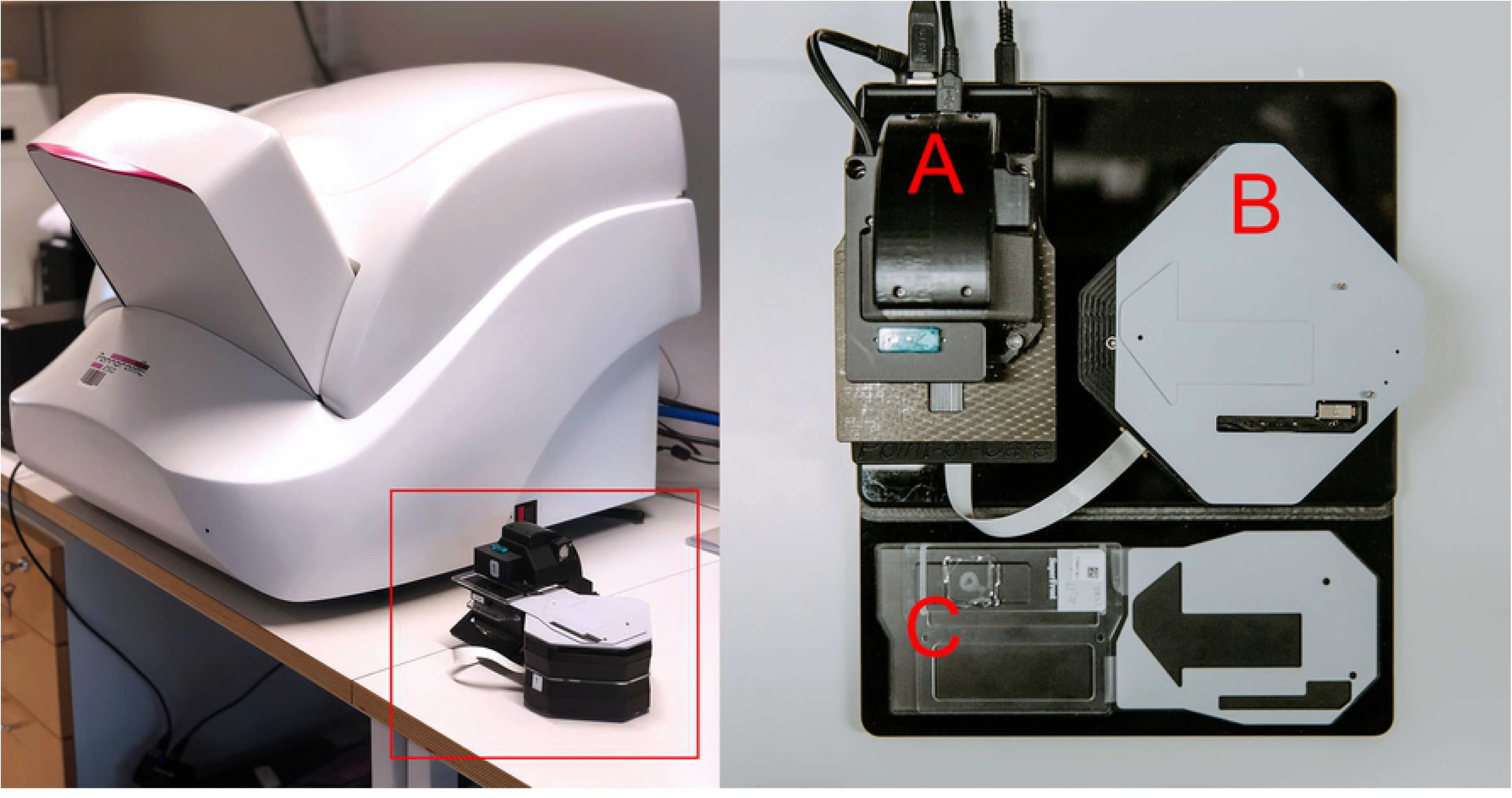
Miniature microscope scanner prototype. Left: Miniature microscope scanner “MoMic” (red bounding box) next to reference whole slide scanner. Right: Overview of the device showing main microscope unit housing camera module (A), motor unit for sample navigation (B) and glass slide holder (C).

**Fig 2.**
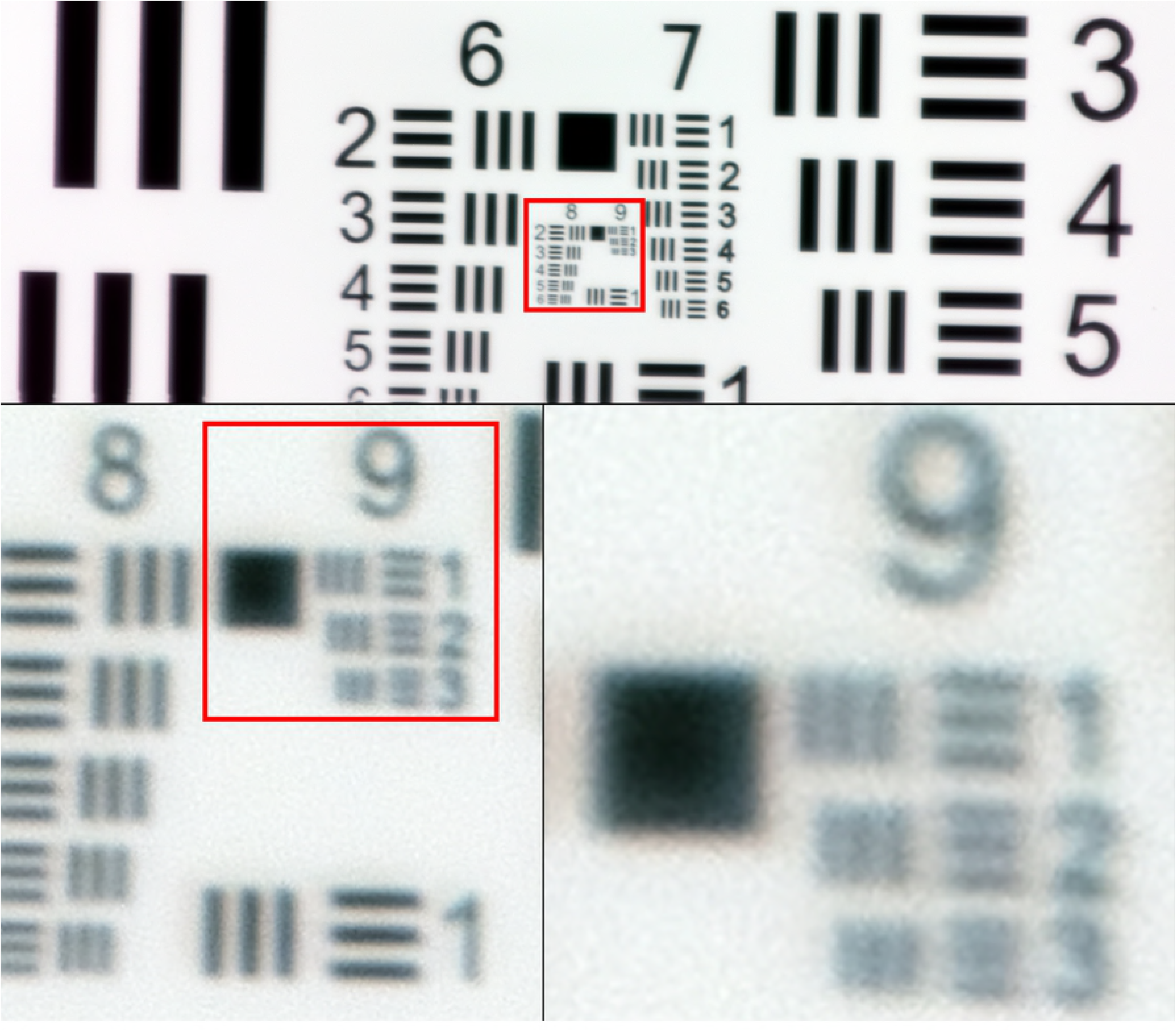
US Air Force 1951 three-bar resolution test chart. Images captured with miniature microscope scanner. Enlarged images showing smallest resolvable bars (group 9, element 2 - 3), corresponding to a spatial resolution of approximately 0.9 µm.

The samples used in this study were also digitized using a high-end, automated whole slide scanner (Pannoramic P250, 3DHistech Ltd., Budapest, Hungary). The slide scanner uses a 20x objective (NA 0.8) equipped with a three-CCD (charge-coupled device) digital camera with a pixel resolution of 0.22 µm. The acquired images were compressed with a compression ratio of 1:9 to a wavelet file format and uploaded to the whole-slide management server, using the configurations described above.

## Slide management and remote analysis of virtual slides

We used the image management platform described above to upload the virtual slides into slide collections for evaluation by the pathologists. Based on these collections of virtual slides, two separate online scoring questionnaires were created for sample evaluations (one for each device) (Fig 3). The scoring system displayed one patient case at a time, starting with the toluidine blue stained sample, after which corresponding anti-cytokeratin stained section was displayed. If only one type of staining was available, only this slide was displayed before continuing to the next case. Display order of patient cases was randomized for both experts, and also for the virtual samples from the separate devices. For every displayed virtual slide, the pathologist was presented with three possible diagnostic categories: “Metastasis”, “No metastasis” and “Not evaluable”. An option for inputting additional comments was also provided for each sample, and experts were encouraged to comment on slide quality during the scoring process. Two independent pathologists evaluated the samples using the online scoring system, which was accessible through a link, sent by e-mail.

**Figure 3.**
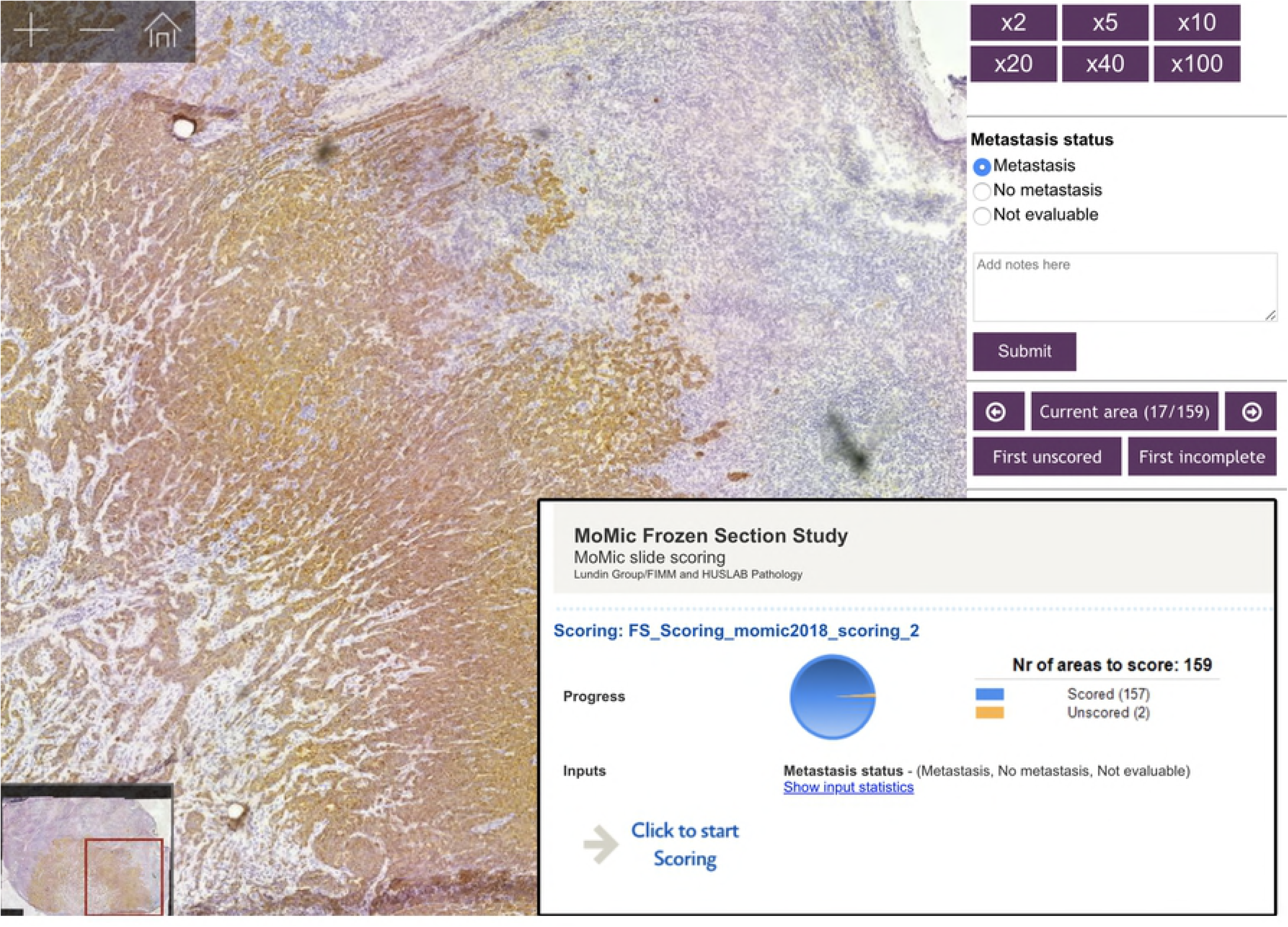
Online image management platform and slide scoring questionnaire. Image showing a scanned lymph node frozen sections (digitized with the miniature microscope scanner) and viewed on the slide management platform with the scoring questionnaire.

### Statistical analysis

Analysis of results and statistical calculations were performed using a general-purpose statistical software package (Stata 15.1 for Mac, Stata Corp., College Station, TX, USA). For statistical analysis, individual samples were classified as either positive for metastatic tissue (i.e. presence of either macro- or micrometastases) or negative for metastatic tissue (i.e. no visible cancer cells). Samples not evaluable according to the pathologists were excluded. Concordance between the miniature microscope scanner, the reference slide scanner and the ground truth was estimated with kappa statistics (kappa values 0.01–0.20 were considered as slight, 0.21–0.40 fair, 0.41–0.60 moderate, 0.61–0.80 good and 0.81– 1.00 as high agreement) [20]. Sensitivity for detection of metastatic cells was calculated as the percentage of true positives (TP) divided by TP and false negatives (FN), with conventional light microscopy analysis as the ground truth (GT). Specificity was calculated as the percentage of true negatives (TN) divided by TN and false positives (FP).

## Results

The total number of slides analyzed after exclusion of samples not evaluable by the pathologists was 152, from 79 patients. By light microscopy 27 (34 %) of these patients were node positive, with 24 (30 %) having macrometastases and 3 (4 %) micrometastases. Correspondingly, 52 patients (66 %) were node negative. When comparing analysis of whole slide images (WSIs) scanned with the miniature microscope scanner to the ground truth, mean overall sensitivity for detection of metastases was 90.56 % (93.88 % and 87.23 %), and mean specificity 99.03 % (100.00% and 98.06 %), on a slide level. When comparing analysis of WSIs from the reference slide scanner to the ground truth, a mean sensitivity of 93.80 % (95.92 % and 91.67 %) for detection of metastases was observed and a mean specificity of 99.03 % (98.06 % and 100.00 %) (Table 1).

**Table 1.** Accuracy for detection of metastases by pathologist analysis of virtual slides, scanned with both microscope scanners.

| **Device** | **Sensitivity (%)** | **Specificity (%)** | **False Negative** | **False Positive** | **Not Evaluable** |
| --- | --- | --- | --- | --- | --- |
| **MoMic (Expert 1)** | 93.88 | 100 | 3 | 0 | 2 |
| **MoMic (Expert 2)** | 87.23 | 98.06 | 6 | 2 | 2 |
| **Reference Scanner (Expert 1)** | 95.92 | 98.06 | 2 | 1 | 1 |
| **Reference Scanner (Expert 2)** | 91.67 | 100.00 | 4 | 0 | 3 |

Table showing overall sensitivity, specificity, total number of FN and FP, and slides classified as not evaluable. Results calculated on a slide level, compared to ground truth.

When measuring agreement between experts, a strong concordance between results from the miniature microscope scanner and ground truth was observed for both experts (k = 0.95 and k = 0.87). Results from the analysis of reference scanner WSIs also showed a strong concordance with the ground truth for both experts (k = 0.95 and k = 0.94). Furthermore, strong intraobserver agreement for both pathologists was observed when comparing results from both scanners for the same expert (k = 0.94 and k = 0.92) (Table 2).

**Table 2.**
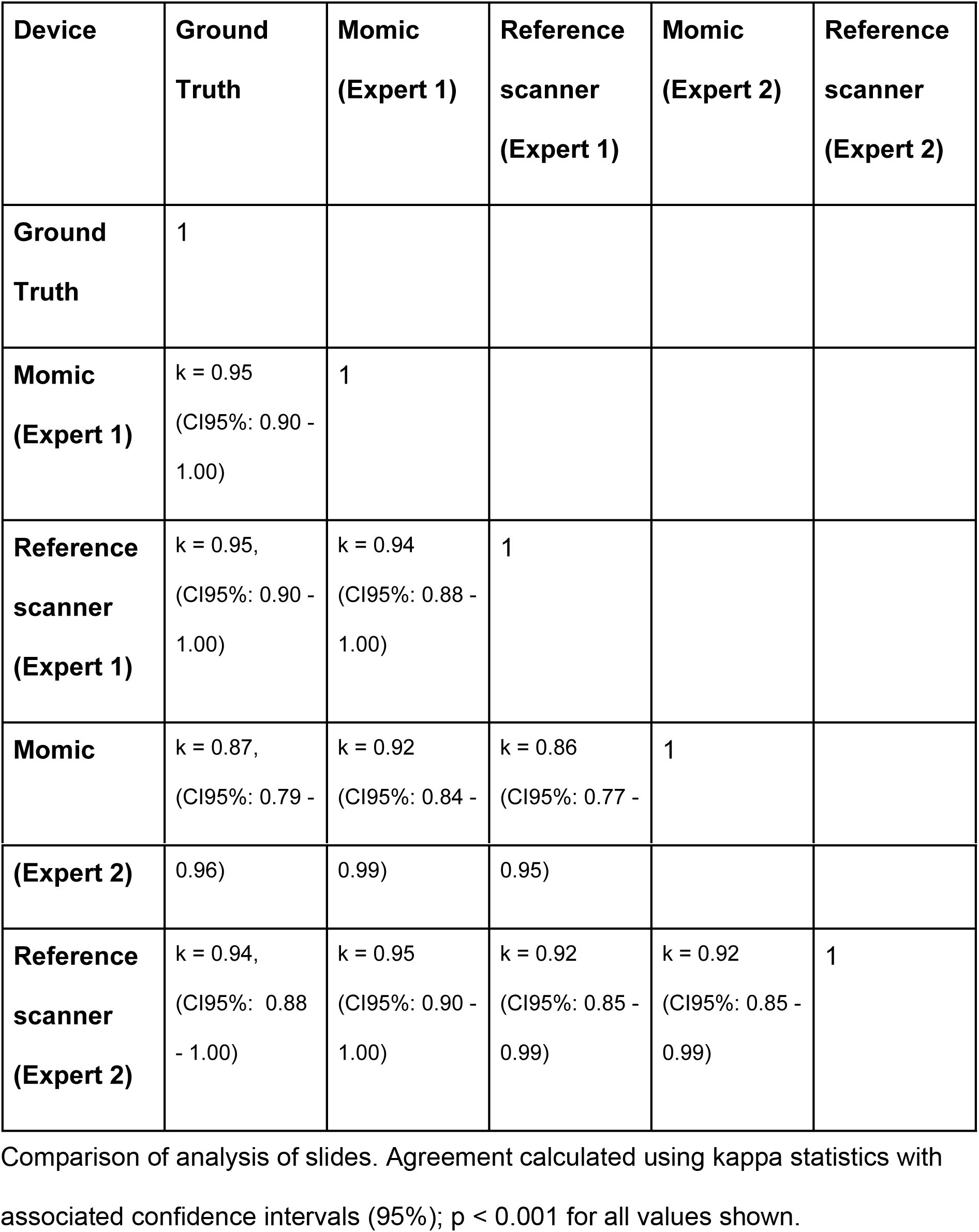
Agreement of results for detection of metastases in slides scanned with the miniature microscope scanner and the reference slide scanner.

The number of false negatives (FN) in the analysis of WSIs scanned with the miniature microscope scanner was 3 (2 %) and 6 (4 %), for the experts. Two samples (1 %) were incorrectly classified as tumor positive by one expert with the miniature microscope scanner, and no false positives (FP) were detected by the other expert. The number of FN slides in the analysis of WSIs from the reference slide scanner was 2 (1 %) and 4 (3 %). For the WSIs from this device, a single FP (1 %) was detected by the first expert, and none by the second. Two slides (1 %) were classified as not evaluable by both experts when analyzing slides from the miniature microscope scanner (different slides for both experts). The number of reference scanner WSIs classified as not evaluable was 1 (1 %) and 3 (2 %).

On a patient level, i.e. including available slides with both staining methods for each patient, the pathologists classified the WSIs from two patients (3 %) incorrectly as tumor negative with the miniature microscope scanner. For the slides scanned with the reference slide scanner, one patient (1 %) was incorrectly classified as tumor negative. This case was the same patient, classified incorrectly as tumor negative in the WSIs from the miniature microscope scanner (Table 3). There were no FP on a patient level with either device. For one patient, both available slides (1 %) were classified as not evaluable by one expert with the miniature microscope scanner.

**Table 3.**
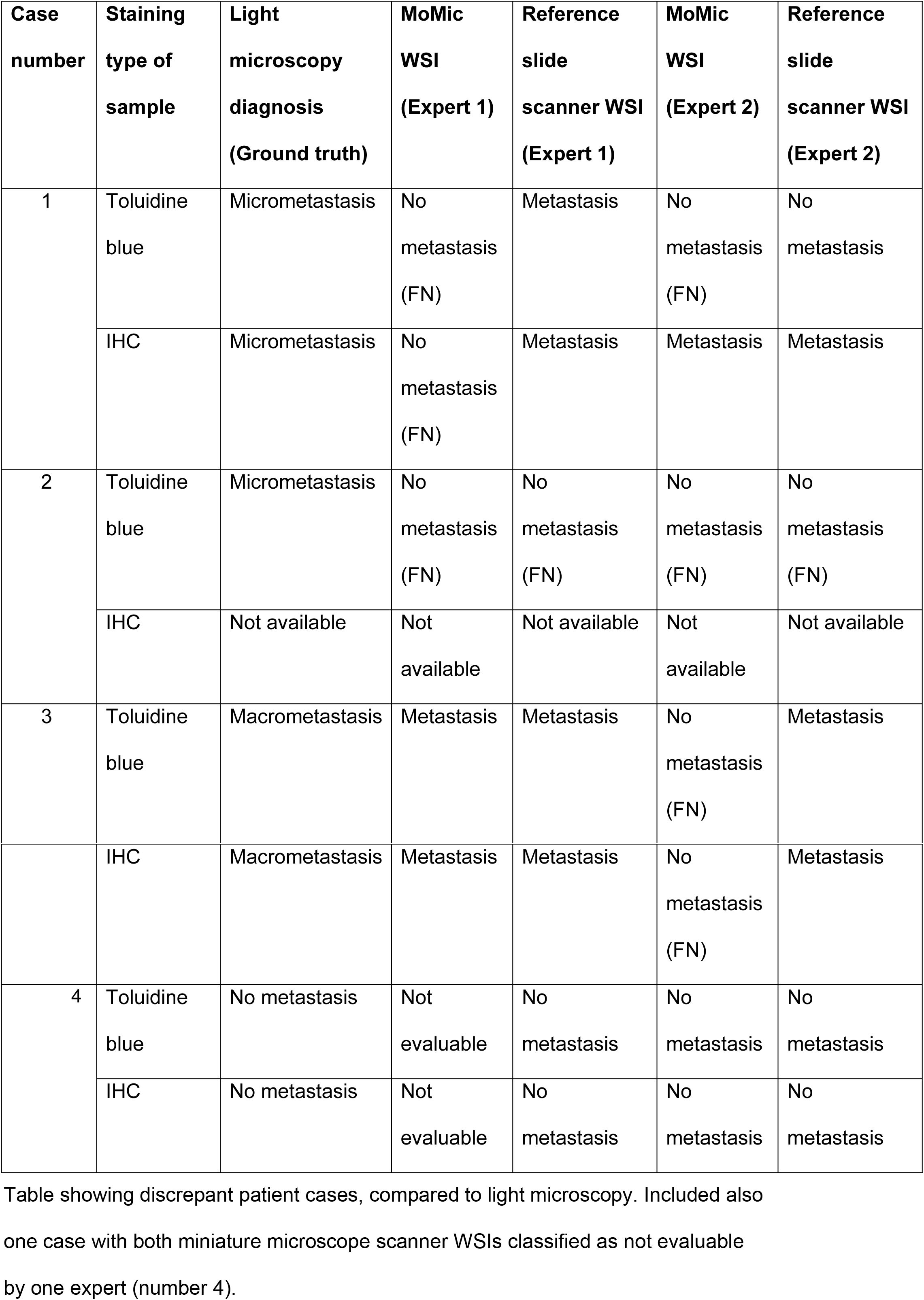
Patient cases diagnosed incorrectly in analysis of WSIs from both scanners, compared to light microscopy.

## Discussion

In this article we evaluate a prototype of a portable, miniature digital microscope scanner for diagnostic assessment of lymph node frozen sections, obtained during breast cancer surgery. Key features of the device include support for whole slide scanning, cloud-connectivity and the use of significantly more inexpensive components, compared to conventional devices. We used the device to digitize archived sentinel lymph node frozen sections and two pathologists with experience in breast cancer histopathology assessed the whole slide images for the detection of metastases. Results were compared to analysis of the same samples scanned with a high-end slide microscope scanner and to pathologist light microscopy analysis of the slides.

Overall, we observed a strong concordance in results from both devices for detection of metastases, compared to light microscopy as study ground truth (GT). A slightly higher concordance to the GT was observed in results from the reference slide scanner (mean *k* = 0.95), than in results from the miniature microscope scanner (mean *k* = 0.91). Agreement between the pathologists was strong (k = 0.92 – 0.94).

Overall sensitivity and specificity for detection of metastases was high for both the miniature microscope scanner (sensitivity 90.56 % and specificity 99.03 %) and the reference slide scanner (sensitivity 93.80 % and specificity 99.03%). Overall, the rate of false negatives (FN) and false positives (FP) was low for both devices, although FN rate was marginally higher with the miniature microscope scanner, and few whole slide images (WSIs) were classified as not evaluable. These results suggest that analysis of slides scanned with both devices yield comparable results, with an overall high grade of sensitivity and specificity.

When grouping available slides from the same patient together, few major discrepancies was observed on a patient level. The slides for two patients were classified incorrectly as negative with the miniature microscope scanner. One of these cases was the same for both experts and also classified incorrectly as negative with the reference slide scanner. This patient had micrometastases, but only toluidine blue-stained sections available, representing a challenging sample (Fig 4). The second patient diagnosed incorrectly as negative by one expert with the miniature microscope scanner also had micrometastases (Fig 5), but both staining methods available. These slides were correctly diagnosed by the second expert. The final discrepant patient case with the miniature microscope scanner, classified incorrectly as negative by one expert, represented a sample with a macrometastasis covering almost the entire section (Fig 6). This sample had sections with both staining methods available, and slides for this case were correctly diagnosed by the second expert. On a patient level, we observed no FP with either device.

**Figure 4.**
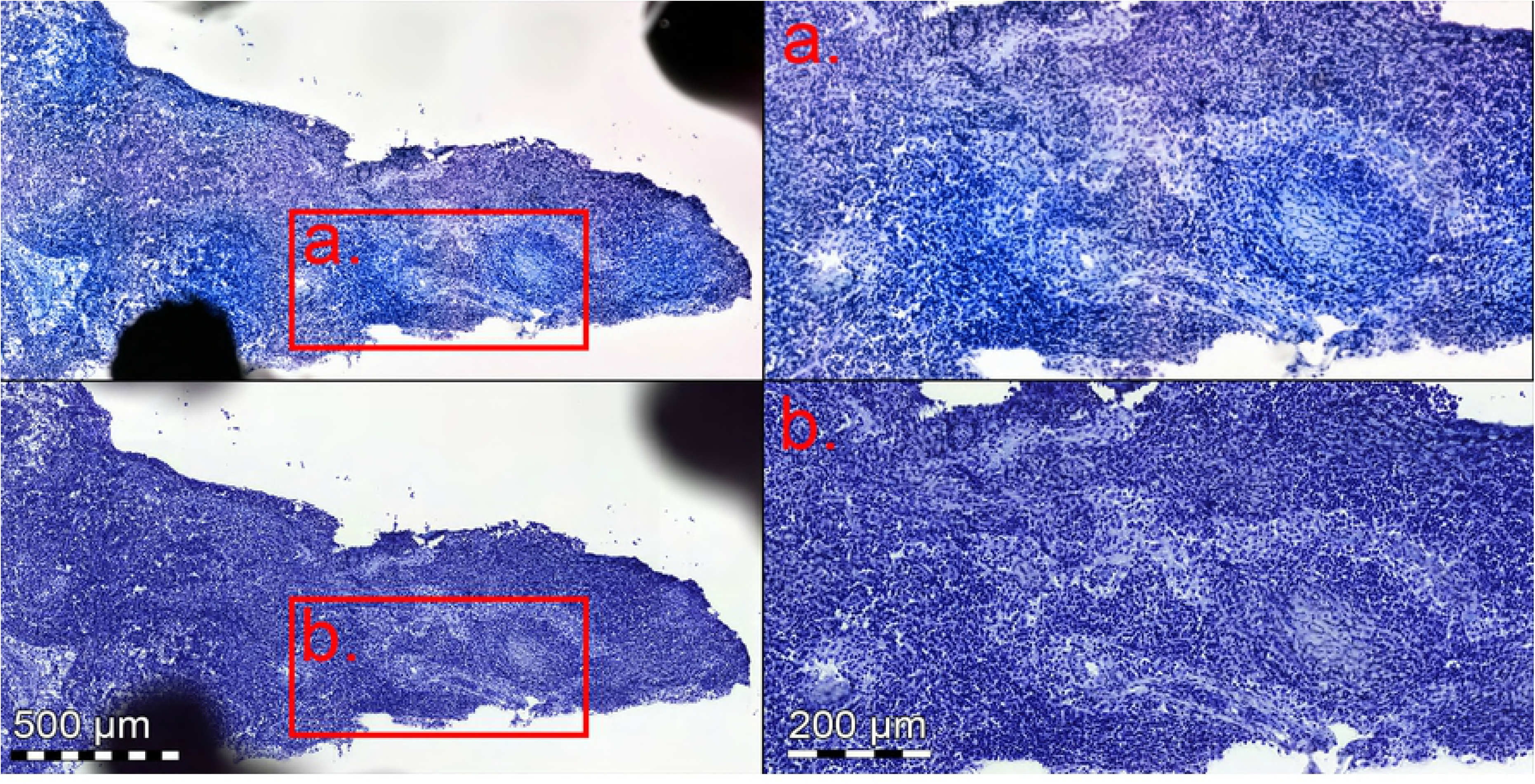
Toluidine blue stained frozen sections with micrometastasis. Slide scanned with the miniature microscope scanner (upper images), and reference slide scanner (lower images). Anti-cytokeratin staining was not available for this section, making analysis challenging, and sample was incorrectly classified as negative by both experts, regardless of device used for digitization.

**Figure 5.**
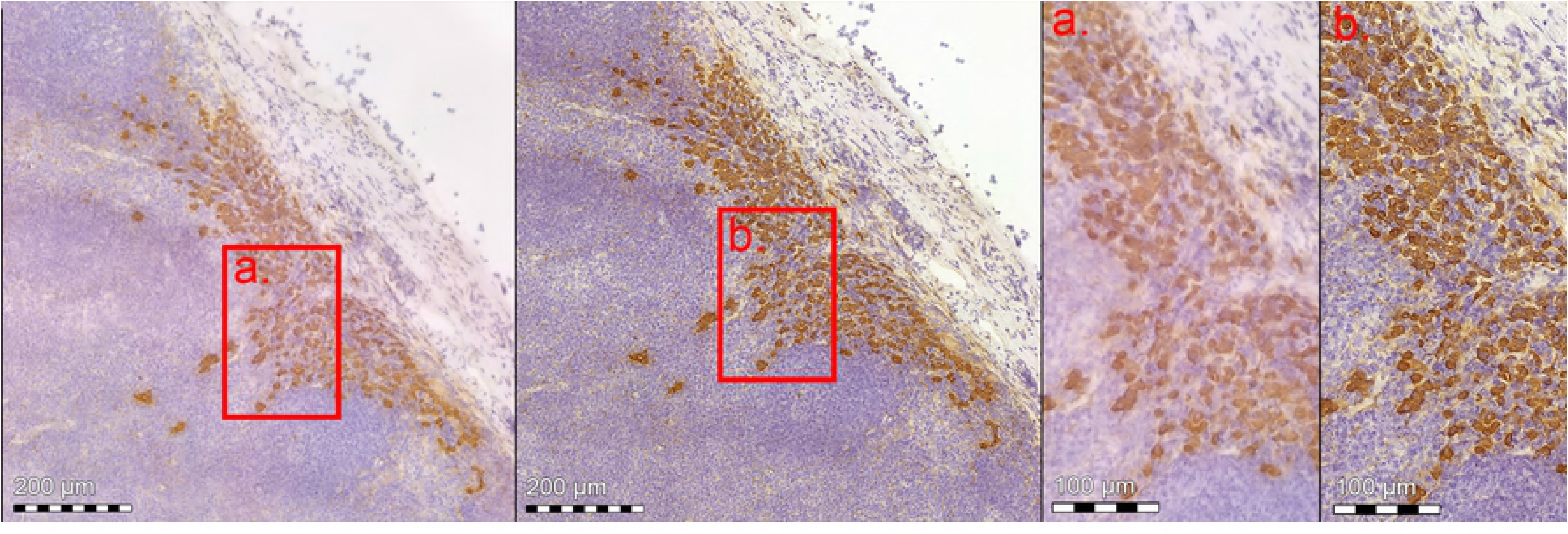
Anti-cytokeratin stained frozen section with micrometastasis. Slide scanned with miniature microscope scanner (left), and reference slide scanner (right). Red bounding box showing higher magnification (a. miniature scanner, and b. reference slide scanner).

**Figure 6.**
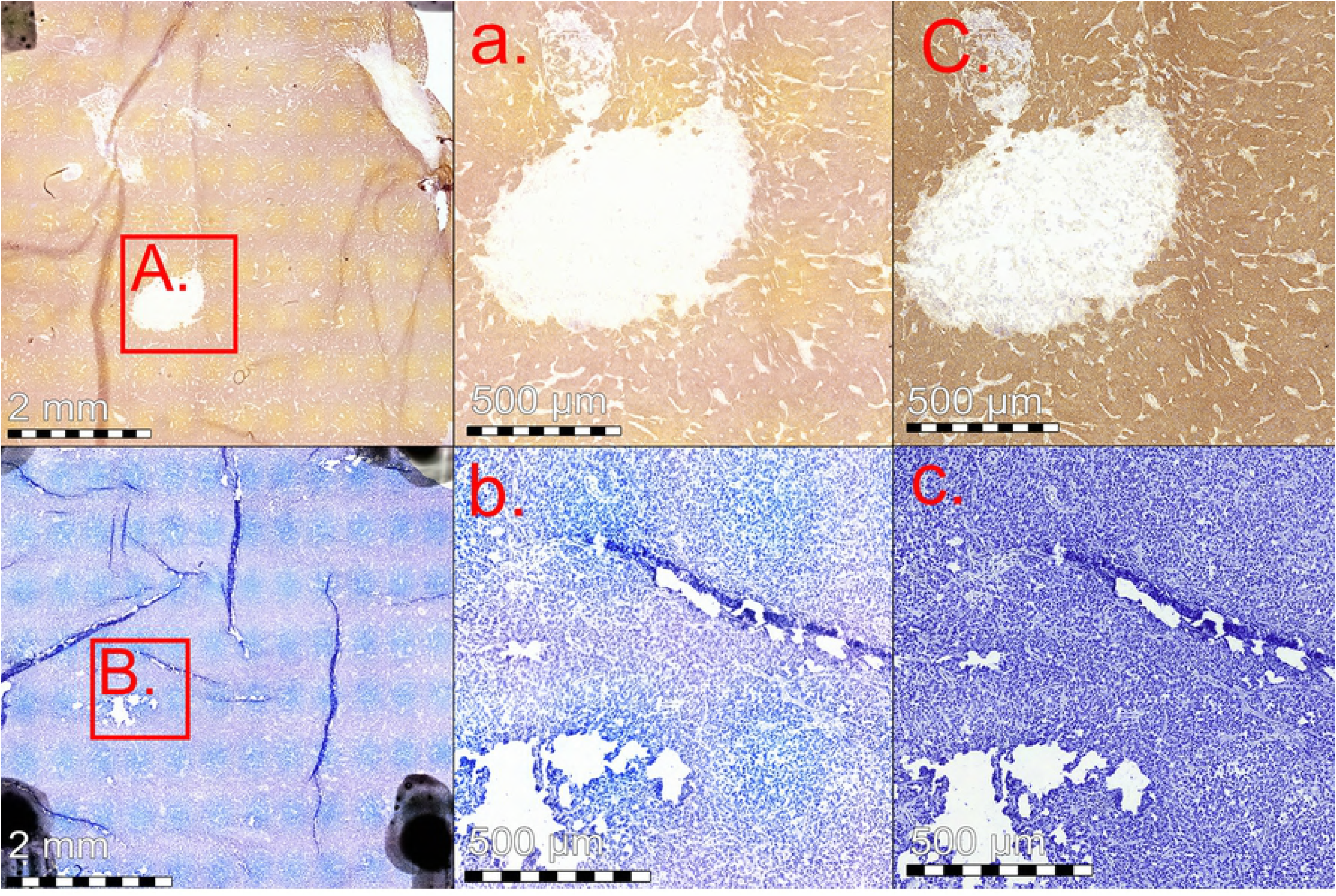
Lymph node frozen section with macrometastasis, stained with both anti-cytokeratin (upper row) and toluidine blue (lower row) staining. Overview of area scanned with miniature microscope (left), red bounding box (A. and B.) showing enlarged area (a. and b.). Reference scanner corresponding regions for comparison purposes shown to the right (C. and c.).

On a slide level, a majority of incorrectly classified WSIs from both devices were toluidine blue slides, and slides with micrometastases. A majority of toluidine blue sections were correctly diagnosed in consecutive anti-cytokeratin stained slides. Detection of metastases in toluidine blue staining is known to be less reliable, especially for smaller lesions [21]. Furthermore, micrometastases in regional lymph nodes are associated with a reduced overall survival, but the exact clinical significance is being debated [22]. Only one major discrepancy was observed on a patient level, were a macrometastasis was misdiagnosed with the miniature microscope scanner WSIs by one expert, suggesting an overall high sensitivity for detection of macrometastases in slides from this device. Among the results from the same expert, a significant part of FN (20 - 30 %) were incorrectly classified in WSIs from both scanners, suggesting other causes for discrepancy than only difference in image quality.

Both pathologists were asked to comment on the quality of virtual slides. A majority of the feedback concerned technical problems, such as areas being out of focus (for both devices), “vignetting” of certain samples (mainly the miniature microscope images) and poor tissue sample quality (in physical slide) (S1 Fig). Most of these problems were related to slide scanning, and could be solved by rescanning affected slides. Interestingly, technical quality did not seem to correlate with accuracy of final sample interpretation, as all samples with comments regarding quality were nonetheless correctly interpreted. Additional comparison WSIs from both devices can be found in the supplementary material (S2 Fig.).

As this in an early study, there is a number of limitations. Our patient material included a relatively small number of micrometastases, and no cases of ITCs. Furthermore, only a single area per slide was digitized. A potential source of bias in this study is that one of the experts analyzing the WSIs (SN) had originally selected the slide areas to be digitized. As WSIs were displayed in a randomized order during analysis, and sample collection was performed in early stages of the study, we believe the risk of significant bias is relatively small. In this study we have focused mainly on image quality of virtual slides, but especially if larger amounts of slides are scanned, additional factors needed to be considered in clinical applications include e.g. turnover time for slide scanning and data connectivity for uploading of digitized slides.

Results here suggest that by using a portable, miniature microscope scanner constructed out of components that are several orders of magnitude cheaper compared to components in currently available scanners, routine breast cancer lymph node frozen sections can be scanned with sufficient quality for detection of metastases. Our work here demonstrates how inexpensive, mass-produced components can be utilized to develop novel solutions for point-of-care slide digitization, and potentially improve access to digital diagnostics and facilitate sample analysis. This technology could likely be expanded also for real time analysis of samples at the point-of-care, e.g. for intraoperative applications. Recent studies show promising results for detection of metastases using deep learning-based image analysis in sentinel lymph node samples, scanned with high-end scanners (23)(24). As our results here suggest that image quality achievable with low-cost components can be comparable, this type of image analysis could likely be applied to samples scanned using this technology also. Further research is needed to validate these results and should focus on evaluating the technology in clinical environments.

## Conclusion

Breast cancer lymph node metastases in frozen sections can be accurately detected by visual analysis of digitized slides, scanned with a low-cost, point-of-care slide scanner, with results comparable to conventional light microscopy and analysis of slides scanned with a high-end whole slide scanner. This method could potentially provide a novel platform for digital diagnostics, especially in resource-limited settings, facilitate sample analysis and reduce the need for experts on-site during surgical procedures.

## Supporting information

**S1 Fig. Technical problems encountered in sample digitization**. A minority of slides scanned with the miniature microscope scanner displayed a “vignetting” effect, showing uneven color intensity in scanned images (left panel). This was likely caused by problems with the brightfield correction algorithm. Areas in some WSIs were out of focus (right panel), due to focusing problems. This affected small areas in a minority of samples, scanned with both devices. Problems here were temporary, and could be corrected by rescanning affected slides.

**S2 Fig. Lymph node frozen section with macrometastasis, stained with toluidine blue (left) and anti-cytokeratin (right) staining, and scanned with both devices**. Upper images showing overview of the FS section and lower side showing enlarged areas (as indicated with red bounding boxes). Slides scanned with the miniature microscope scanner on left side (A. and C.), and reference slide scanner WSIs on the right side for comparison (B. and D.).

## References

(1) Siegel RL, Miller KD, Jemal A. Cancer statistics, 2018. CA: a Cancer Journal for Clinicians 2018 Jan;68(1):7-30.

(2) Fisher B, Bauer M, Wickerham DL, Redmond CK, Fisher ER, Cruz AB, et al. Relation of number of positive axillary nodes to the prognosis of patients with primary breast cancer. An NSABP update. Cancer 1983 Nov 01;52(9):1551-1557.

(3) Mansel RE, Fallowfield L, Kissin M, Goyal A, Newcombe RG, Dixon JM, et al. Randomized multicenter trial of sentinel node biopsy versus standard axillary treatment in operable breast cancer: the ALMANAC Trial. J Natl Cancer Inst 2006 May 03;98(9):599-609.

(4) Samphao S, Eremin JM, El-Sheemy M, Eremin O. Management of the axilla in women with breast cancer: current clinical practice and a new selective targeted approach. Annals of Surgical Oncology 2008 May;15(5):1282-1296.

(5) Weiser MR, Montgomery LL, Susnik B, Tan LK, Borgen PI, Cody HS. Is routine intraoperative frozen-section examination of sentinel lymph nodes in breast cancer worthwhile?. Annals of Surgical Oncology 2000 Oct;7(9):651-655.

(6) Lucci A, McCall LM, Beitsch PD, Whitworth PW, Reintgen DS, Blumencranz PW, et al. Surgical complications associated with sentinel lymph node dissection (SLND) plus axillary lymph node dissection compared with SLND alone in the American College of Surgeons Oncology Group Trial Z0011. Journal of Clinical Oncology 2007 Aug 20;25(24):3657-3663.

(7) Liu L, Lang JE, Lu Y, Roe D, Hwang SE, Ewing CA, et al. Intraoperative frozen section analysis of sentinel lymph nodes in breast cancer patients. Cancer 2011;117(2):250-258.

(8) Holck S, Galatius H, Engel U, Wagner F, Hoffmann J. False-negative frozen section of sentinel lymph node biopsy for breast cancer. Breast 2004 Feb;13(1):42-48.

(9) Yeh Y, Nitadori J, Kadota K, Yoshizawa A, Rekhtman N, Moreira AL, et al. Using frozen section to identify histological patterns in stage I lung adenocarcinoma of <= 3 cm: accuracy and interobserver agreement. Histopathology 2015 Jun;66(7):922-938.

(10) Weinstein RS, Descour MR, Liang C, Barker G, Scott KM, Richter L, et al. An array microscope for ultrarapid virtual slide processing and telepathology. Design, fabrication, and validation study. Hum Pathol 2004 Nov;35(11):1303-1314.

(11) Al-Janabi S, Huisman A, Van Diest PJ. Digital pathology: current status and future perspectives. Histopathology 2012 Jul;61(1):1-9.

(12) Gifford AJ, Colebatch AJ, Litkouhi S, Hersch F, Warzecha W, Snook K, et al. Remote frozen section examination of breast sentinel lymph nodes by telepathology. ANZ J Surg 2012 Nov;82(11):803-808.

(13) Thorstenson S, Molin J, Lundström C. Implementation of large-scale routine diagnostics using whole slide imaging in Sweden: Digital pathology experiences 2006-2013. Journal of Pathology Informatics 2014;5(1):14.

(14) Isaacs M, Lennerz JK, Yates S, Clermont W, Rossi J, Pfeifer JD. Implementation of whole slide imaging in surgical pathology: A value added approach. Journal of Pathology Informatics 2011;2:39.

(15) Zhu H, Isikman SO, Mudanyali O, Greenbaum A, Ozcan A. Optical imaging techniques for point-of-care diagnostics. Lab on a Chip 2013 Jan 07;13(1):51-67.

(16) Holmström O, Linder N, Ngasala B, Mårtensson A, Linder E, Lundin M, et al. Point-of-care mobile digital microscopy and deep learning for the detection of soil-transmitted helminths and Schistosoma haematobium. Glob Health Action 2017 Jun;10(sup3):1337325.

(17) Pirnstill CW, Cote GL. Malaria Diagnosis Using a Mobile Phone Polarized Microscope. Scientific Reports 2015 Aug 25;5:13368.

(18) Holmström O, Linder N, Lundin M, Moilanen H, Suutala A, Turkki R, et al. Quantification of Estrogen Receptor-Alpha Expression in Human Breast Carcinomas With a Miniaturized, Low-Cost Digital Microscope: A Comparison with a High-End Whole Slide-Scanner. PLoS ONE [Electronic Resource] 2015;10(12):e0144688.

(19) Konsti J, Lundin M, Linder N, Haglund C, Blomqvist C, Nevanlinna H, et al. Effect of image compression and scaling on automated scoring of immunohistochemical stainings and segmentation of tumor epithelium. Diagnostic Pathology 2012;7:29.

(20) Landis JR, Koch GG. The measurement of observer agreement for categorical data. Biometrics 1977 Mar;33(1):159-174.

(21) Chao C. The use of frozen section and immunohistochemistry for sentinel lymph node biopsy in breast cancer. Am Surg 2004 May;70(5):414-419.

(22) Andersson Y, Bergkvist L, Frisell J, de Boniface J. Long-term breast cancer survival in relation to the metastatic tumor burden in axillary lymph nodes. Breast Cancer Research & Treatment 2018 Sep;171(2):359-369.

(23) Ehteshami Bejnordi B, Veta M, Johannes van Diest P, van Ginneken B, Karssemeijer N, Litjens G, et al. Diagnostic Assessment of Deep Learning Algorithms for Detection of Lymph Node Metastases in Women With Breast Cancer. JAMA 2017;318(22):2199-2210.

(24) Liu Y, Kohlberger T, Norouzi M, Dahl GE, Smith JL, Mohtashamian A, et al. Artificial Intelligence–Based Breast Cancer Nodal Metastasis Detection. Arch Pathol Lab Med 2018.

